# Motion-Dependent Object Perception Reveals Limits of Current Video Neural Networks

**DOI:** 10.64898/2026.03.15.711964

**Authors:** Matteo Dunnhofer, Jean de dieu Uwisengeyimana, Kohitij Kar

## Abstract

How does motion contribute to robust object perception when appearance cues are unreliable? In natural scenes, camouflage, clutter, and occlusion can obscure object boundaries in static images, yet humans often resolve these ambiguities once objects move. Here we ask whether modern artificial vision systems capture this motion-dependent computation and whether their internal representations align with those used by biological vision. Using videos from the MOCA (Moving Camouflaged Animals) dataset, we introduce behavioral benchmarks that quantify the accuracy of object position and size estimation in scenes containing either static or moving objects. We first compare human observers with a diverse set of artificial neural networks. For static stimuli, image-based and video-based models achieve similar accuracy in predicting object position and size. However, humans show systematic improvements when objects are presented in motion. Image-based models do not exhibit this motion-dependent improvement, whereas several video-based architectures reproduce this behavioral pattern by integrating information across time. To examine the representational basis of these differences, we record neural population responses from the macaque inferior temporal (IT) cortex during presentation of the same stimuli. Models that more closely match IT representations also better reproduce human motion-dependent behavior. These results show that static accuracy alone is insufficient to evaluate models of visual perception and that alignment with primate visual representations provides a useful guide for developing models that capture dynamic computations in vision.

## 1 Introduction

How does motion contribute to robust object perception when appearance cues are unreliable? In natural environments, objects are frequently embedded in clutter, partially occluded, or concealed by camouflage, making their boundaries difficult to infer from static images alone. Yet humans and other animals often resolve such ambiguities once objects move. Motion can reveal structure that is otherwise indistinguishable from the background, suggesting that dynamic information plays an important role in stabilizing visual perception.

Modern computer vision systems, however, are largely built on static image recognition. Deep convolutional neural networks trained on large-scale image classification datasets have achieved remarkable performance and now form the foundation of many visual recognition systems (Krizhevsky et al. 2012; He et al. 2016; Dosovitskiy et al. 2020). The internal layers of these models also partially predict neural responses in the primate ventral visual stream that houses key circuits to solve object recognition (Yamins et al. 2014; Cadena et al. 2019; Bashivan et al. 2019; Schrimpf et al. 2018), leading them to be interpreted as the current best computational hypotheses for the representations observed in primates (Kar and DiCarlo 2024). However, because these architectures process frames independently, they cannot explicitly exploit temporal structure present in dynamic scenes, and therefore do not align with primates under many scenarios (Yakubovskaya et al. 2026; Dunnhofer et al. 2026a).

In contrast, humans appear to benefit substantially from motion when perceiving objects in challenging environments. Camouflaged animals, for example, can remain nearly invisible when stationary but become detectable when they move (Fig. 1 A). Behavioral studies (Julesz 1971; Braddick 1993) suggest that dynamic information improves object detection and localization when appearance cues are weak or ambiguous. These observations raise an important question: do modern artificial vision systems capture the motion-dependent computations that support robust object perception in biological vision?

**Fig. 1.**
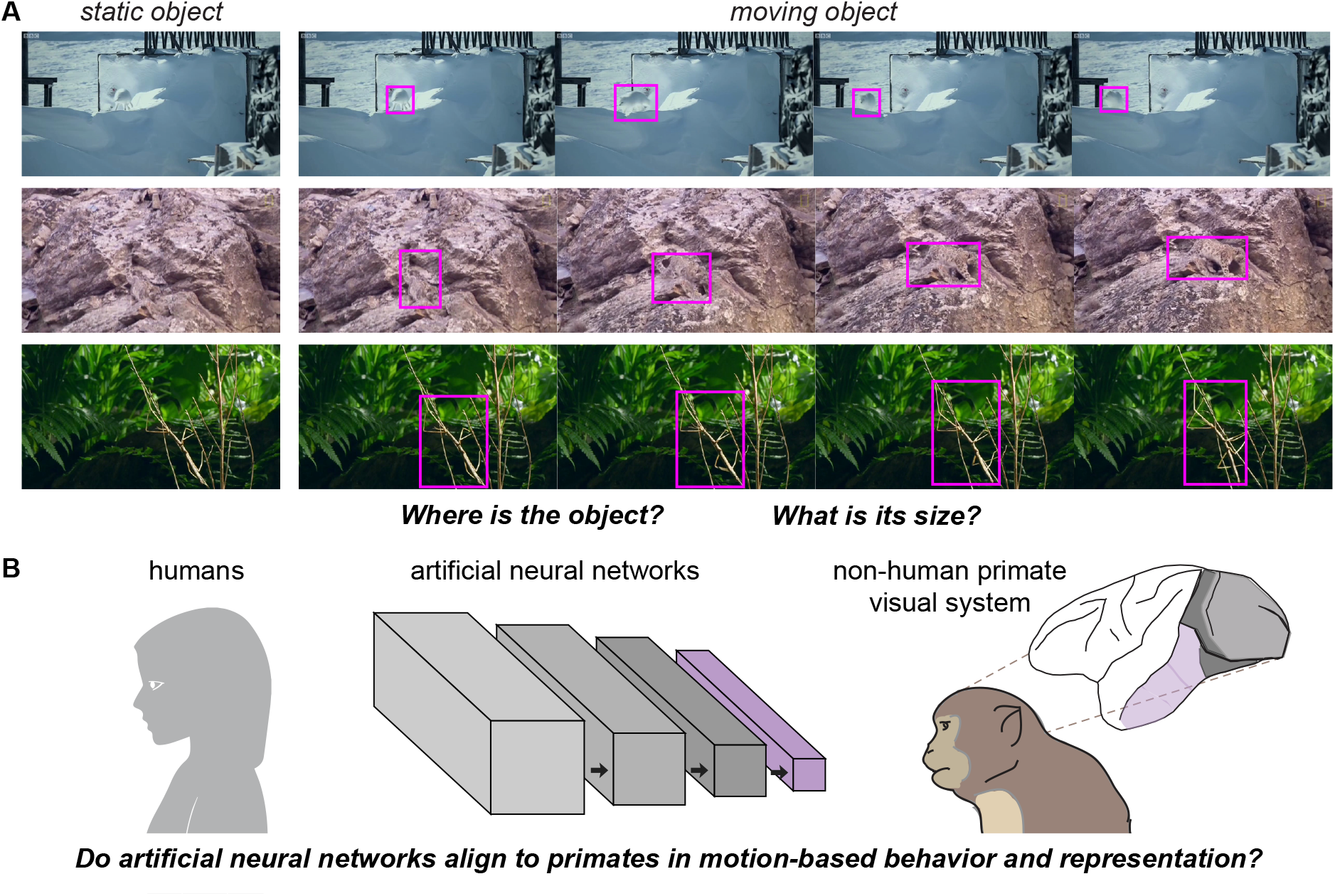
Motion-dependent object perception across humans, artificial neural networks, and the primate visual system. A. Examples of camouflaged animals shown as static images and as frames from short (500 ms) videos. When objects remain static they are difficult to detect, but motion reveals their structure and location. Purple boxes indicate ground-truth object annotations used for behavioral and decoding analyses. B. Overview of the experimental framework. We compare three systems: human observers, artificial neural networks (ANNs), and neural population responses in macaque inferior temporal (IT) cortex, on the same task of estimating object attributes under camouflage. This unified framework allows us to evaluate both behavioral alignment between humans and models and representational alignment between models and the primate visual system.

Biological visual systems are well suited to exploit such information. Classical theories distinguish between ventral and dorsal visual pathways specialized for object recognition and motion processing respectively (Goodale and Milner 1992). While this framework has been highly influential, increasing evidence suggests that ventral visual areas also encode dynamic information. Neurons in inferotemporal (IT) cortex represent not only object identity but also attributes such as position, size, and transformations of visual stimuli (Logothetis et al. 1995; Hong et al. 2016). Recent work further indicates that temporal dynamics can enhance the reliability and informativeness of population codes in IT (Ramezanpour et al. 2024; Dunnhofer et al. 2026a,b). These findings suggest that temporal information may contribute directly to stabilizing object representations when appearance cues are degraded.

Despite these insights from neuroscience and perception, most computational models of object recognition remain fundamentally static. Image-based neural networks process each frame independently and therefore cannot exploit temporal structure. In contrast, a growing class of video-based neural network architectures integrates information across time using spatiotemporal operations (Tran et al. 2015; Carreira and Zisserman 2017; Bertasius et al. 2021). These models achieve strong performance on video understanding tasks, yet it remains unclear whether their representations capture the motion-dependent computations that support robust object perception in biological systems.

In this work, we investigate whether motion improves the fidelity of object form estimation and whether modern artificial vision systems reproduce this effect. To address this question, we introduce behavioral benchmarks based on the MOCA (Moving Camouflaged Animals) dataset (Lamdouar et al. 2020). These benchmarks quantify the accuracy with which humans or models estimate object attributes such as position and size in scenes containing either static or moving objects. By comparing performance on static and dynamic versions of the same stimuli, we isolate the contribution of motion to object perception.

We evaluate these benchmarks across three levels of analysis (Fig. 1 B). First, we measure behavioral performance in human observers performing object localization and size estimation tasks. Second, we evaluate a diverse set of artificial neural networks (ANNs), comparing image-based models that process frames independently with video-based architectures that integrate temporal information across video frames. Third, we analyze neural population responses recorded from macaque inferotemporal cortex while animals view the same stimuli. This unified framework allows us to directly compare behavioral performance, computational models, and neural representations of dynamic visual information.

Our results reveal three key findings. First, human observers show systematic improvements in estimating object position and size when motion information is available. Second, image-based neural networks achieve strong static performance but do not exhibit this motion-dependent improvement. Third, several video-based architectures reproduce the behavioral pattern, and the degree to which they do so is predicted by their representational similarity to macaque IT population responses.

Together, these findings suggest that static object accuracy alone is insufficient to evaluate models of visual perception. Instead, the ability to reproduce motion-dependent improvements observed in human behavior provides a stronger test of model validity. Our results further indicate that alignment with primate visual representations can serve as a useful guide for identifying artificial models that capture dynamic computations underlying natural vision.

### Contributions

This work makes four main contributions:

- **Benchmark for motion-based object perception**. We introduce new behavioral benchmarks based on the MOCA dataset that quantify how accurately humans and models estimate object position and size in challenging camouflaged scenes where appearance cues alone are unreliable.
- **Evidence that motion stabilizes object form perception**. We show that human observers systematically improve their estimation of object position when motion cues are available, and that neural population responses in macaque inferior temporal (IT) cortex exhibit a corresponding improvement in the fidelity of object attribute representations.
- **Evaluation of artificial neural networks on motion-based object perception**. We demonstrate that image-based neural networks can accurately estimate object form attributes under camouflage but fail to exploit motion cues in a primate-like way, whereas video-based architectures that integrate temporal information show human-like motion-dependent improvements in decoding object attributes.
- **Brain-guided evaluation of artificial models**. By comparing human behavior, neural population responses, and artificial neural networks within a unified framework, we show that models whose internal representations more closely align with macaque IT cortex better predict human perceptual patterns, although important gaps between biological and artificial systems remain.

## 2 Related Work

Our work lies at the intersection of motion perception, robust object recognition, and biologically inspired computer vision. We review prior work in five areas: motion-based object perception, motion-defined form and structure-from-motion, video-based neural network architectures, camouflage and challenging natural scenes, and neural studies of motion and form interactions.

### 2.1 Motion as a cue for object perception

Motion is one of the most powerful cues for separating objects from background in natural scenes. Classic work in visual perception demonstrated that coherent motion can reveal object structure even when static appearance cues are weak or ambiguous (Julesz 1971; Braddick 1993). Differences in motion between foreground and background provide a strong signal for perceptual grouping and figure–ground segmentation (Rauber and Treue 1998; Braddick and Atkinson 2011).

In computer vision, motion cues have long been used to detect and segment objects in videos. Early approaches relied on optical flow and motion boundaries to identify moving regions (Horn and Schunck 1981; Shi et al. 1994). Later work integrated motion cues into tracking and video segmentation frameworks (Brox and Malik 2010; Wang and Schmid 2013). More recently, deep learning approaches have leveraged motion information for tasks such as video object segmentation and tracking (Tokmakov et al. 2017; Yan et al. 2021; Oh et al. 2019; Dunnhofer et al. 2023, 2025).

Despite these advances, most object recognition systems still operate primarily on static appearance cues (Krizhevsky et al. 2012; He et al. 2016; Dosovitskiy et al. 2020), leaving the computational role of motion in stabilizing object form largely unexplored.

### 2.2 Motion-defined form and structure-from-motion

A closely related line of research investigates how motion alone can reveal object structure. In structure-from-motion paradigms, three-dimensional shape can be recovered from the motion of points projected onto the retina or image plane (Ullman 1979). Psychophysical studies have demonstrated that observers can recover detailed object structure even when individual frames contain minimal shape information (Treue et al. 1995). These findings highlight the remarkable ability of biological vision to infer form from dynamic signals.

In computer vision, structure-from-motion techniques have traditionally been used to reconstruct 3D geometry from image sequences (Tomasi and Kanade 1992; Agarwal et al. 2011; Wang et al. 2025). However, most work in this area focuses on geometric reconstruction rather than object recognition. As a result, relatively little attention has been devoted to understanding how motion-derived signals contribute to the perception of object attributes such as position, scale, or shape under challenging visual conditions.

### 2.3 Video-based artificial neural networks

Recent progress in deep learning has produced a wide range of neural network architectures designed to process video data. Early work extended image-based convolutional networks into the temporal domain using 3D convolutions (Tran et al. 2015). Subsequent models introduced more powerful spatiotemporal architectures such as Inflated 3D ConvNets (Carreira and Zisserman 2017), SlowFast networks (Feichtenhofer et al. 2019), and transformer-based video models (Bertasius et al. 2021; Arnab et al. 2021; Li et al. 2024). These models have achieved strong performance on tasks such as action recognition and video classification.

While these architectures explicitly integrate temporal information, they are typically optimized for action recognition rather than object perception. Consequently, it remains unclear whether their temporal representations capture the same motion-based mechanisms that support robust object perception in biological systems. In particular, the potential role of temporal integration in stabilizing object form representations under challenging visual conditions has received relatively little attention.

### 2.4 Camouflage and challenging natural scenes

Camouflage represents a particularly challenging setting for visual recognition systems. When an object’s appearance closely matches its background, static appearance cues become unreliable. In such cases, motion often provides the most reliable signal for detecting and localizing objects. Recent work in computer vision has begun to investigate detection and segmentation of camouflaged objects (Fan et al. 2020; Mei et al. 2021; Lamdouar et al. 2023). Several datasets have been proposed to evaluate recognition performance under these conditions. Among them, the MOCA (Moving Camouflaged Animals) dataset provides naturalistic videos in which animals are often indistinguishable from their background in static frames but become visible through motion (Lamdouar et al. 2020).

While prior work has largely focused on segmentation and detection, less attention has been devoted to understanding how motion improves the estimation of object attributes such as position or size.

### 2.5 Neural studies of motion and form interactions

A large body of neuroscience research has examined how motion and form are represented in the primate visual system. Classical theories of visual processing propose a functional division between ventral and dorsal pathways, with the ventral stream specialized for object recognition and the dorsal stream specialized for motion and spatial processing (Goodale and Milner 1992). Under this framework, motion processing is typically associated with dorsal areas such as MT and MST.

However, increasing evidence suggests that ventral visual areas also encode dynamic information (Bigelow et al. 2023; Ramezanpour et al. 2024). Neurons in the IT cortex represent not only object identity but also attributes such as position, size, and transformations of visual stimuli (Hong et al. 2016). Functional imaging studies further show that ventral areas can respond to motion-defined form stimuli (Vanduffel et al. 2002).

More recent work has begun to investigate the role of temporal dynamics in shaping ventral representations. Studies comparing neural responses to static images and videos suggest that dynamic stimuli can enhance the reliability and informativeness of IT population codes (Ramezanpour et al. 2024; Dunnhofer et al. 2026b,a). At the same time, goal-driven modeling has demonstrated that deep neural networks trained on object recognition tasks can predict neural responses in ventral visual cortex with surprising accuracy (Yamins et al. 2014; Cadena et al. 2019; Schrimpf et al. 2018). Most of these models, however, operate on static images. Whether modern video-based neural networks better capture the temporal computations performed by ventral cortex remains largely unexplored.

### 2.6 Summary

Taken together, prior work suggests that motion can play a powerful role in object perception. However, three key gaps remain. First, the computational contribution of motion to stabilizing object form representations under naturalistic camouflage has not been systematically quantified. Second, it remains unclear whether modern video-based artificial neural networks capture the same motion-based advantages observed in biological vision. Third, few studies have directly compared behavioral performance, neural population codes, and artificial models within a unified framework.

## 3 Methods

### 3.1 Ethics Approval

#### Human Experiments

The study was approved by the York University Ethics Review Committee (Human Participant Review Subcommittee). Participants were recruited via the Amazon Mechanical Turk platform. Consent was presented on the first page; only consenting participants proceeded. Compensation rate was 15 CAD/hour. **Non human primate experiments:** All data were collected, and animal procedures were performed, in accordance with the NIH guidelines, the Massachusetts Institute of Technology Committee on Animal Care, and the guidelines of the Canadian Council on Animal Care on the use of laboratory animals and were also approved by the York University Animal Care Committee.

### 3.2 Visual stimuli

All experiments in our study were conducted using videos from the MOCA dataset (Lamdouar et al. 2020). This dataset contains naturalistic video clips of animals embedded in complex environments where object appearance closely matches the background, creating strong camouflage (see Fig. 1 A for exemplar frames).

From the dataset we selected 132 video clips containing camouflaged animals moving within their natural environment. Each clip was truncated to a duration of 500 ms (30 frames at 60 Hz) for presentation in behavioral and neural experiments. In addition to the moving videos, a static object control condition was created by extracting the first frame from the same videos and presenting them without motion.

Bounding-box-based ground-truth annotations provided with the dataset were used to extract object attributes including horizontal position (*x*), vertical position (*y*) (barycenter of bounding-box), object size (area of bounding-box), and object speed in the pixel space (mean barycenter shift of bounding-box). These annotations served as the target variables for both behavioral analyses and model decoding.

### 3.3 Human behavioral experiments

Human observers (*N* = 154) performed perceptual estimation tasks using the aforementioned video stimuli via Amazon Mechanical Turk. Each trial began with a fixation period of 300 ms followed by a 500 ms presentation of a video clip depicting either a moving or static object. After the stimulus presentation, participants were instructed to report object attributes using an interactive interface. Two behavioral tasks were performed:

- **Object localization**. Participants clicked on the perceived center of the object within the scene.
- **Object size estimation**. Participants adjusted a bounding-box to match the perceived size of the object.

Participants were given up to 5000 ms to complete each response. Behavioral accuracy was measured by computing the absolute pixel error between participant responses and ground-truth object annotations. Visual illustration of the trials and behavioral tasks are given in Fig. 2 A-B. To assess the benefit of motion, performance was compared between trials containing videos with moving objects and trials containing videos with static objects.

**Fig. 2.**
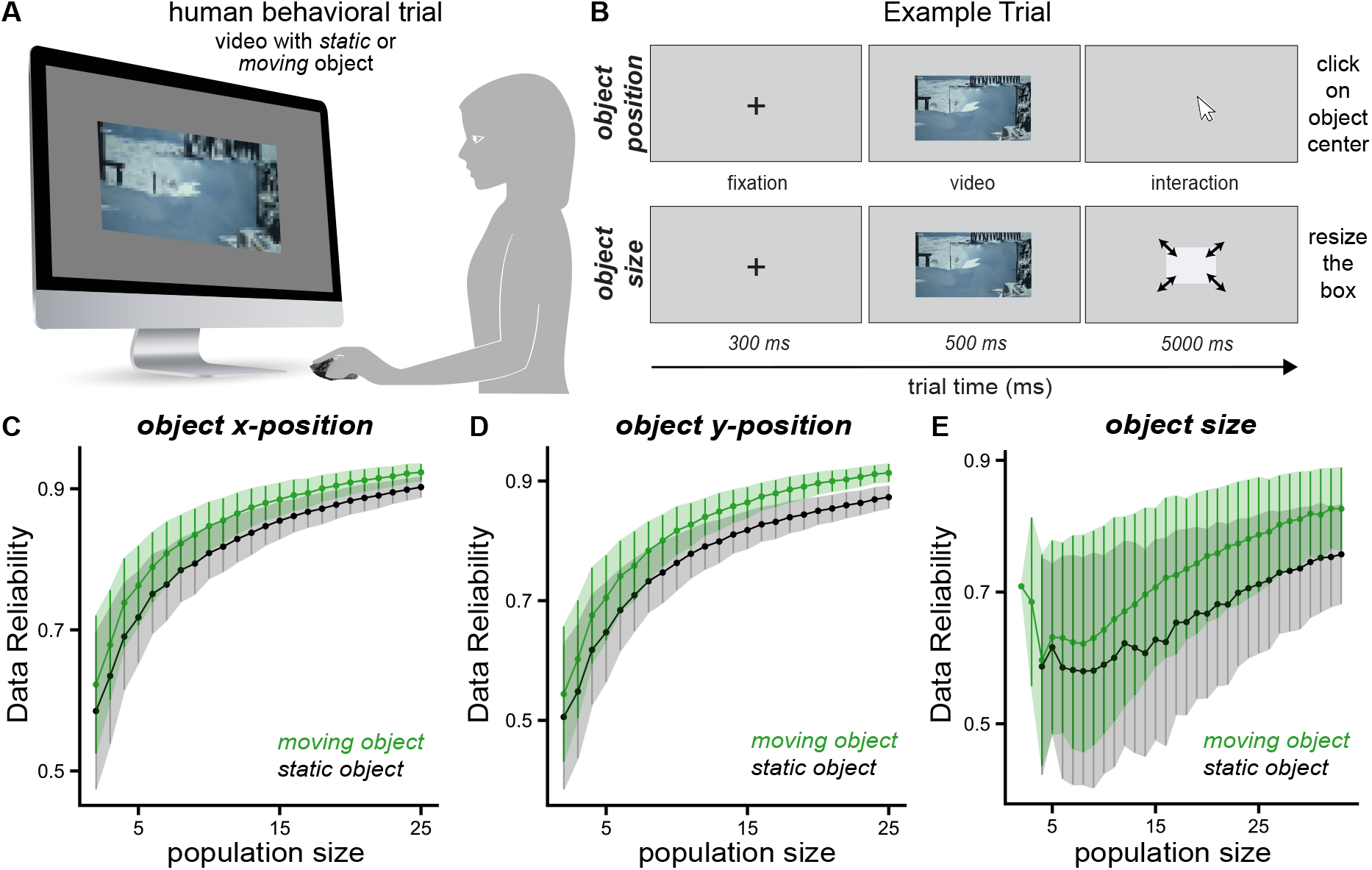
Human behavioral tasks and reliability of object attribute estimates. A. Example behavioral trial. Participants fixated for 300 ms, viewed a 500 ms video containing either a static or moving camouflaged object, and then reported either the object center (localization task) or object size (bounding-box adjustment) within a time limit of 5000 ms. B. Schematic of the two behavioral tasks used in the study: object localization and object size estimation. C–E. Splithalf reliability of human behavioral estimates for the objects’ horizontal position (C), vertical position (D), and size (E) as a function of the number of repetitions in measurement of each stimuli. Reliability is consistently higher when objects move (green) compared to when they remain static (gray), indicating that motion stabilizes perceptual estimates of object attributes.

### 3.4 Non-human primate electrophysiology

We measured neural responses in two adult rhesus macaques (*Macaca mulatta*). Animals passively viewed the same set of video stimuli while maintaining fixation on a central fixation point. Neural activity was recorded across the inferior temporal (IT) cortex using Utah microelectrode arrays (Blackrock Microsystems). In total, neural activity was recorded from 192 sites from each monkey. After initial fixation for 300 ms, stimuli were presented for 500 ms, in the central 10 degree of visual field of the monkeys while animals maintained fixation. Neural responses were recorded continuously during stimulus presentation. Spike trains were converted into firing-rate responses by computing spike counts within sliding temporal windows aligned to stimulus onset (for details, see Kar et al. (2019)).

### 3.5 Neural decoding of object attributes

To quantify the information about object attributes (position and size) contained in neural population responses, we trained linear decoders to predict the variables from the IT population activity patterns, as visualized in Fig. 3.

**Fig. 3.**
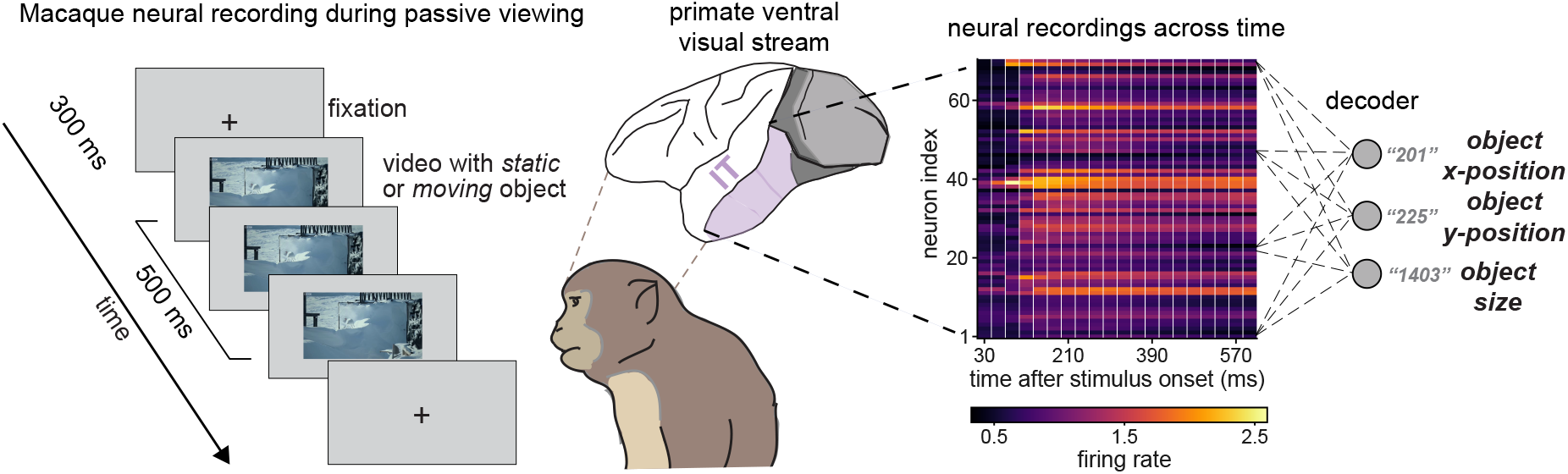
Neural population decoding of object attributes from macaque IT cortex. Monkeys passively viewed the same video stimuli used in the human behavioral experiment while neural activity was recorded from inferior temporal (IT) cortex. Monkeys initiated a trial by fixating on a dot for 300 ms. We then presented the stimuli for 500 ms followed by an inter trial interval of 500 ms. Spike responses were converted to firing rates and used to train linear decoders predicting object horizontal position, vertical position, and size. This analysis quantifies the amount of information about object attributes present in IT population responses and allows direct comparison with human behavior and artificial neural network representations.

For each stimulus presentation, neural responses were represented as a population activity vector across recorded sites. For each time bin, neural firing rates were first averaged across a temporal window extending from each earlier times up to the current bin, yielding stimulus-by-neuron response matrices that capture accumulated activity over time. Regularized linear regression models (either Partial Least Squares or Ridge regression) were then trained to predict the following attributes:

- object horizontal position (*x*)
- object vertical position (*y*)
- object size.

Decoder performance was evaluated using cross-validation across stimuli with 20 folds. Prediction accuracy was measured using Spearman correlation between predicted and ground-truth values. To estimate the reliability of the decoding procedure, we computed a split-half consistency metric. Trials were randomly divided into two non-overlapping halves, and separate regression models were trained on the responses from each half. Predictions obtained from the two halves were then compared by computing the Spearman correlation between them. This procedure provided an estimate of split-half reliability, which was subsequently corrected using the Spearman–Brown formula to account for the reduced number of trials in each half (Yamins et al. 2014; Kar et al. 2019; Hong et al. 2016). Decoding accuracy was defined as the maximum score across time bins. We then compared decoding accuracies obtained from moving-object and static-object stimuli to determine whether object motion improves the fidelity of neural representations of object attributes.

### 3.6 Artificial neural network models

To compare biological and artificial vision systems in response to camouflaged object motion, we evaluated a diverse set of generalistic artificial neural networks for computer vision tasks, including both image-based and video-based architectures, whose details are reported in Table 1.

**Table 1.**
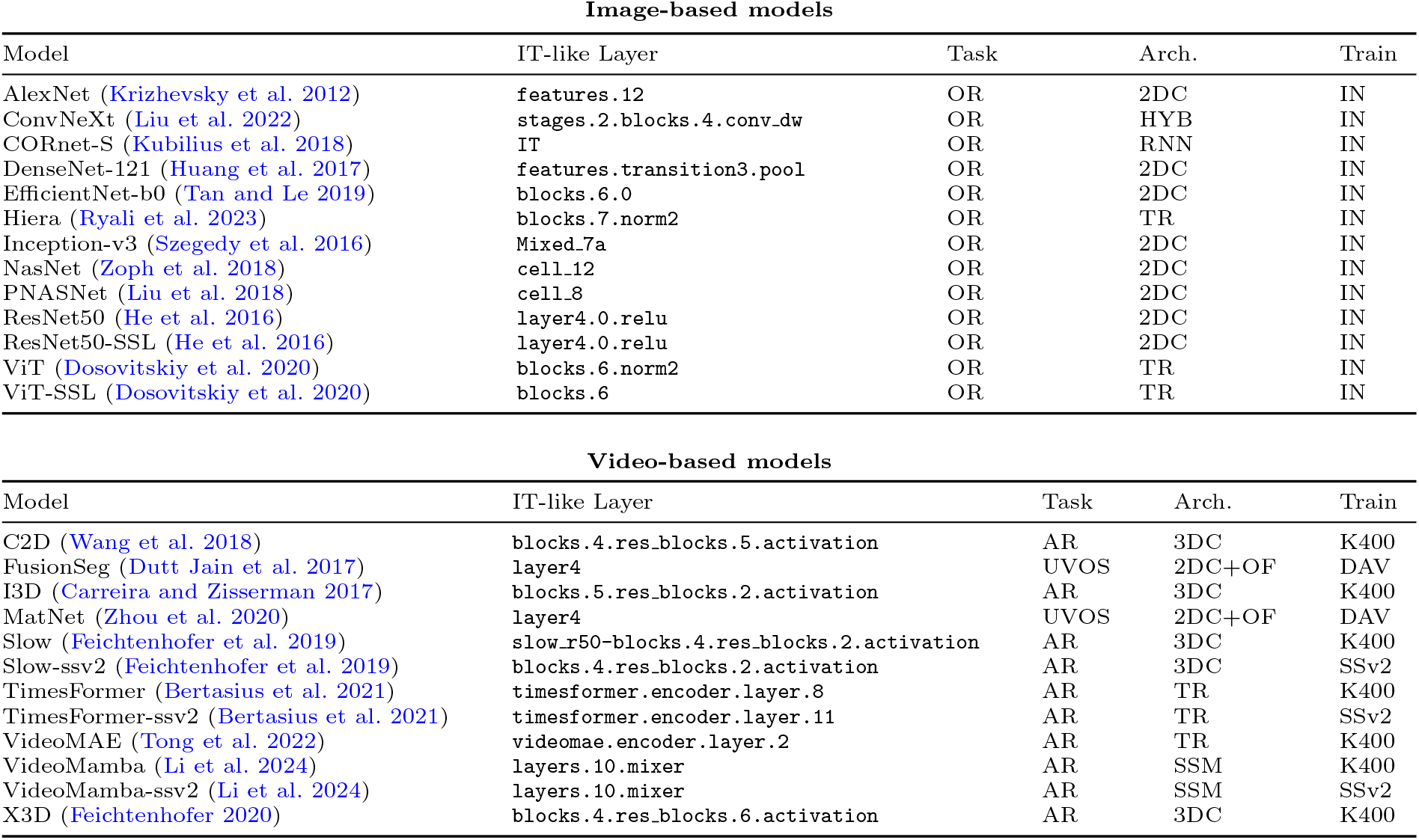
Artificial neural networks architectures tested in this study. We report: architecture name; the corresponding IT-like layer used for feature extraction; the task the architecture was optimized for; type of architecture; training dataset. Task acronyms: OR = Object Recognition, AR = Action Recognition, UVOS = Unsupervised Video Object Segmentation, SVOS = Semi-supervised Video Object Segmentation. Architecture acronyms: 2DC = 2D Convolutional Network, 3DC = 3D Convolutional Network, TR = Transformer, RNN = Recurrent Neural Network, HYB = Hybrid architecture, SSM = State Space Model, 2DC+OF = 2D Convolution with Optical Flow. Training datasets: IN = ImageNet, K400 = Kinetics-400, SSv2 = Something-Something-v2, DAV = DAVIS.

#### Image-based models

These models process individual video frames independently without temporal integration. Frames from each video (composed of 30 frames) were passed through the network separately, and feature activations were extracted from the IT-like feature layer reported on the Brain-Score benchmark (Yamins et al. 2014; Schrimpf et al. 2018).

#### Video-based models

We extracted frame-indexed feature activations from different types of video-based neural networks. Following prior work in spatio-temporal architecture interpretation (Kowal et al. 2022, 2024), we selected models by varying tasks and considering their accuracy and popularity across computer vision benchmarks. For video-based models with a fixed temporal buffer of *F* frames (e.g., action recognition models), features were extracted at each frame in a sliding window approach by filling the buffer with the current frame and the *F −*1 preceding frames in a first-in-first-out (FIFO) manner. When fewer than *F* previous frames were available, the buffer was filled by repeating the first frame. To identify the IT-like layer of each video ANN, we extracted features from multiple evenly spaced layers and tested each feature against macaque IT neural responses using an held-out image-response dataset (Dapello et al. 2023; Sörensen et al. 2024). For each layer, neural predictivity (Yamins et al. 2014; Kar et al. 2019) was computed by predicting neural responses to the same images. The layer achieving the highest percentage of explained variance (%EV) was selected as the IT-like layer of the video model (Schrimpf et al. 2018).

All models used in this study were off-the-shelf, publicly available, pretrained models and hyperparameters from their original repositories. No additional training or fine-tuning was performed. Input transformations were implemented according to the original model repository. For all models, feature representations were compressed to a population of 1000 units using sparse random projection multiple times.

### 3.7 Decoding object attributes from model features

For each model, feature activations were extracted for all stimuli and used to train linear decoders predicting object attributes (position, size, and speed) at each video frame separately. The decoding model training and testing followed the neural analysis pipeline used for IT recordings, and it is visualized in Fig. 4 A-B, separately for image-based and video-based models.

**Fig. 4.**
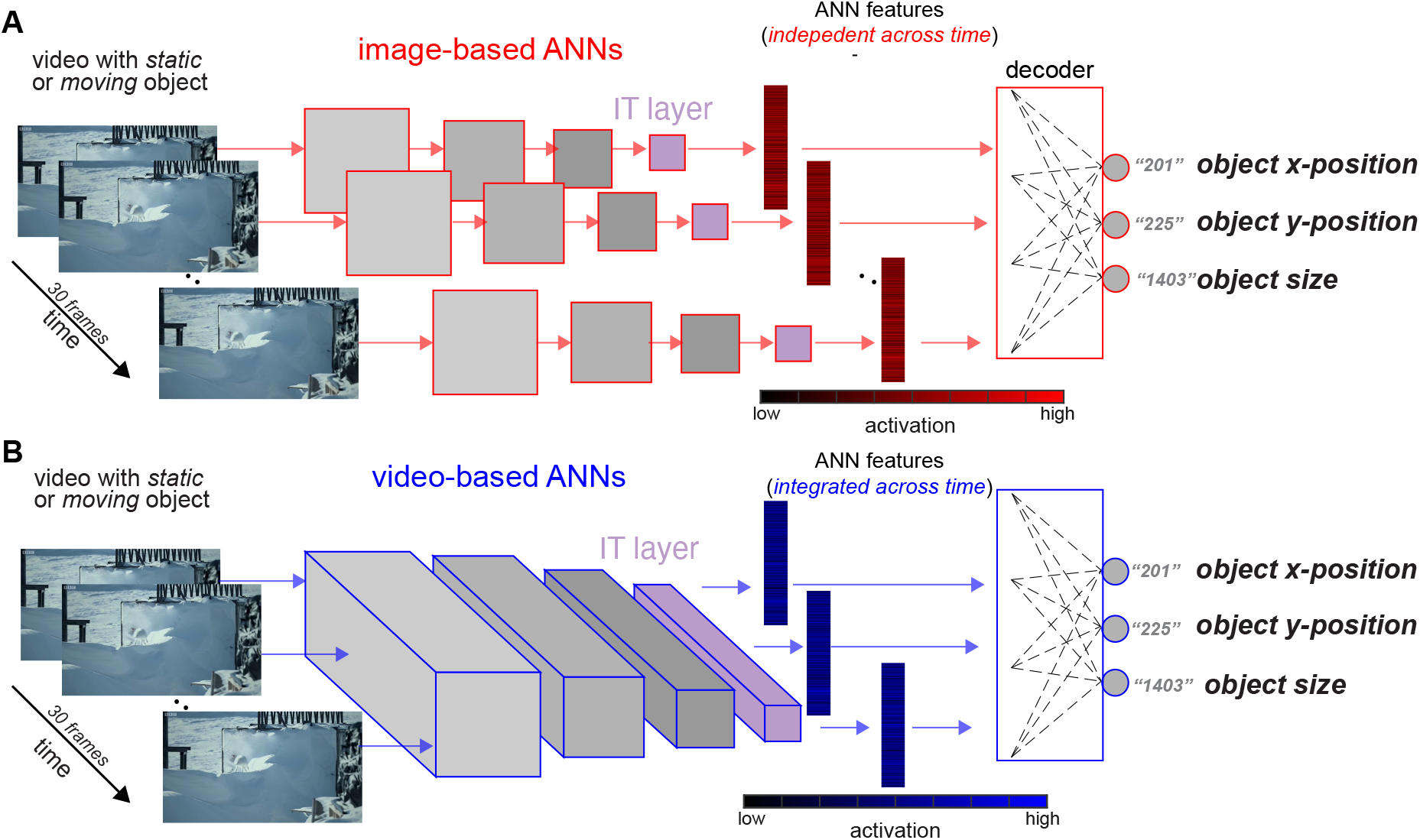
Decoding object attributes from artificial neural network representations. A. Analysis pipeline for image-based neural networks. Each frame of the video stimulus is processed independently, and feature activations are extracted from the model’s IT-like layer. Linear decoders are then trained to estimate object horizontal position, vertical position, and size from these representations. B. Same as in A, but for video-based models. In this case, frame-wise features were extracted using temporal integration of consecutive frames, which were jointly processed by the model through spatiotemporal operations (i.e., instead of individual frames, the video was presented to the ANNs). Decoders were then trained to estimate the object’s horizontal (*x*) and vertical (*y*) position, as well as its size, from these temporally integrated representations. This framework mirrors the neural decoding analysis performed on macaque IT responses, enabling direct comparison between biological and artificial representations.

Model performance was quantified as decoding accuracy by computing the Spearman correlation between predicted and ground-truth object attributes. The retained decoding accuracy was defined as the maximum score across frame-wise decodes. To assess the contribution of motion information, decoding accuracy was compared between moving object videos and static object videos.

### 3.8 Brain-model alignment

To quantify the representational similarity between artificial neural networks representation and neural population responses, we computed Centered Kernel Alignment (CKA) (Kornblith et al. 2019) between model features and neural activity. For each time bin, neural firing rates were aggregated over temporal windows extending from earlier bins up to the current bin, yielding stimulus-by-neuron response matrices that capture the accumulation of neural activity over time. In these analyses, we only retained visually reliable neural recording sites that passed a reliability threshold of 0.4 based on split half trial-to-trial response consistency (78 for monkey and 51 for monkey 2).

In parallel, frame-based representations were extracted from each model for the corresponding video frames. CKA was then computed between the model features and the neural response matrix across stimuli, providing a measure of representational similarity. To account for noise in the neural recordings, we estimated neural reliability using a split-half procedure in which stimulus repetitions were randomly divided into two halves and CKA was computed between the corresponding neural response matrices. Model–neural CKA scores were subsequently normalized using this split-half estimate to obtain noise-corrected similarity values. This analysis was repeated across time bins to characterize how well model representations at each frame align with the temporally evolving neural population responses. For each model, we retained the CKA score relating the model representation and neural time bin achieving the highest human behavior consistency.

### 3.9 Model comparison and alignment

Model performance was compared with human performance and neural decoding results obtained from IT cortex. Scatter plots comparing correlations for static object and moving object stimuli were used to evaluate whether models showed improvements similar to humans and neural responses. Models were grouped into imagebased and video-based categories. Neural decoding performance was plotted alongside model results to assess the degree of alignment between biological and artificial representations. Histograms were used used to compare families of models against humans and neural responses. Barplots were used to compare different models among themselves.

### 3.10 Statistical Analyses

All statistical analyses were performed in Python using the scipy.stats library. Relationships between continuous variables (e.g., human prediction, neural or model decodes) were assessed using Spearman rank correlation coefficients, unless otherwise specified.

Group comparisons followed a consistent procedure based on the distributional properties and pairing of the data. Normality of each distribution was assessed using the Shapiro–Wilk test. When both groups satisfied the normality assumption, parametric tests were applied: a paired t-test for paired comparisons (e.g., within-subject or within-model analyses) and an independent-samples t-test for unpaired comparisons (e.g., between models). If the normality assumption was violated for at least one group, non-parametric tests were used instead. Specifically, we applied the Wilcoxon rank-sum test (equivalent to the Mann–Whitney U test) for unpaired comparisons. All tests were two-tailed, and exact p-values and test statistics are reported.

The analyses in this study involve several conceptually independent comparisons (e.g., model-to-human, model-to-brain, architecture-specific trends). Because these analyses test distinct hypotheses, each was evaluated within its own statistical family rather than pooled across the entire manuscript. Following common practice in computational and systems neuroscience, we therefore report exact p-values and test statistics without applying global multiple-comparison corrections. Results that do not reach statistical significance are described as non-significant trends and are not used to support inferential conclusions.

## 4 Results

### 4.1 Human observers benefit from motion when estimating object attributes under camouflage

Motion provides a powerful cue for segmenting objects from background when appearance information is unreliable. If motion contributes directly to stabilizing object representations, then observers should estimate object attributes more accurately when the same scenes are viewed as dynamic videos rather than static frames. We therefore tested whether motion improves human estimates of object position and size in camouflaged scenes. Human participants viewed short video clips from the MOCA dataset and performed object localization (*n* = 91) and size estimation (*n* = 131) tasks (Fig. 5). To isolate the contribution of motion, we compared performance on dynamic videos with performance on static frames extracted from the same clips, ensuring that the underlying visual content remained identical while temporal information was removed. We observed that when objects moved in the videos, observers exhibited significantly lower localization error compared to static conditions. This improvement was observed for both horizontal (x) position (Δ*error*(*static− moving*) = 8.4 px, *p <* 0.001, *t*(131) = 3.9) and vertical (y) position estimates (Δ*error* = 2.8 px, *p* = 0.01, *t*(131) = 2.5). A qualitatively similar but non significant effect was observed for object size estimation. Participants produced more accurate size estimates when viewing dynamic stimuli relative to static frames (Δ*error*(*static −moving*) = 349.6 px, *p* = 0.69, *t*(131) = 0.39). Interestingly, the magnitude of the improvement in the behavioral estimates increased specifically for the stimuli that were more difficult under static conditions (Fig. B-D), suggesting that motion provides information that helps disambiguate objects that are otherwise difficult to detect due to camouflage. In Fig. 5 B-D), the analysis was performed in a cross validated way to avoid correlational artifacts. The stimuli choices in the x-axis were estimated from a different set of subjects and the Δ*error* was estimated on a held-out set of subjects for the static condition. Consistent with these findings, we also observed that the trial split half reliability of object form (position, size) estimates, when measured for the movies with object motion, was significantly higher than when estimated for the corresponding static images (see Fig 2).

**Fig. 5.**
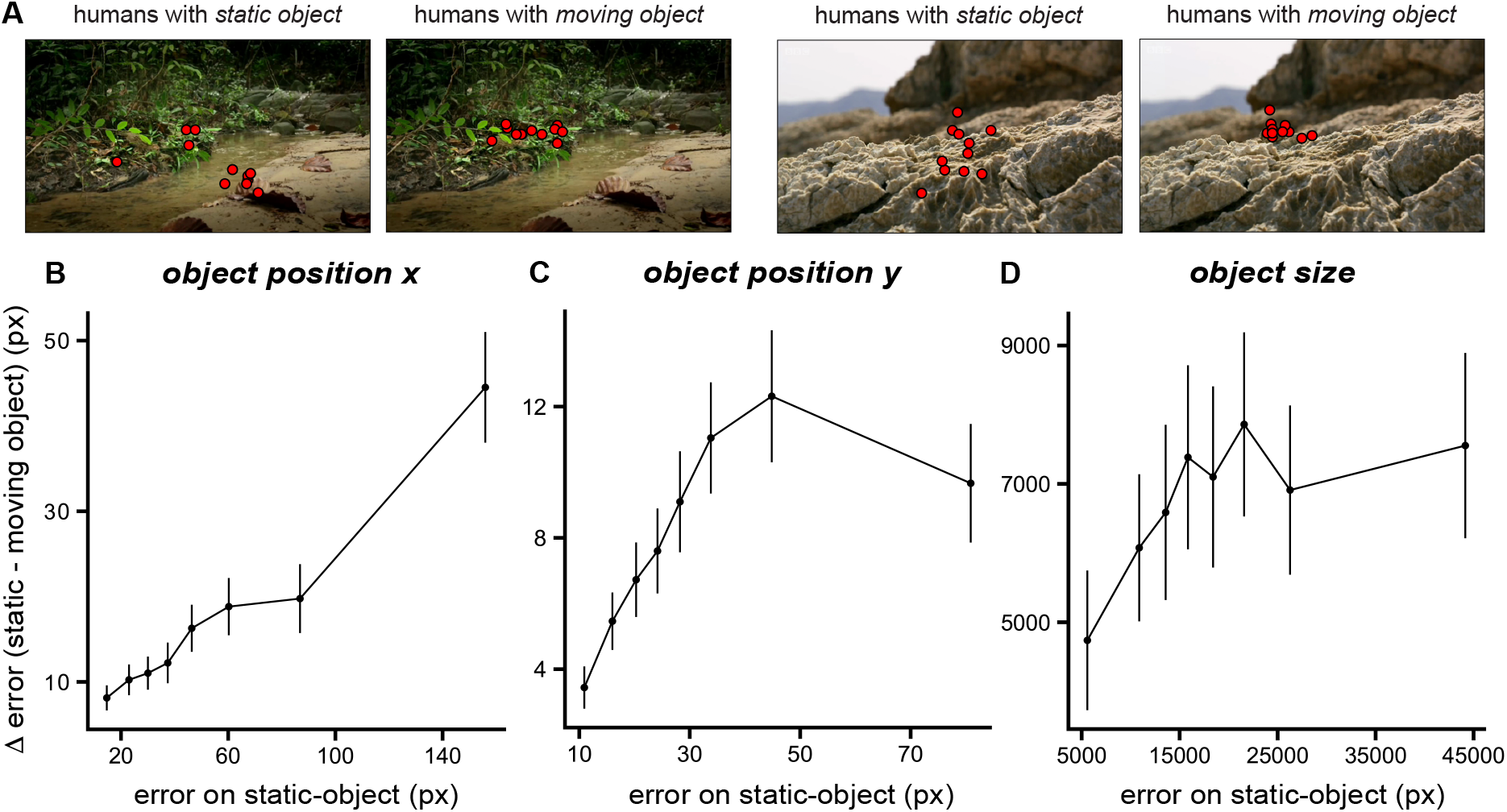
Motion improves human estimation of object position under camouflage. A. Example behavioral responses when observers viewed static versus moving objects. Motion produces more consistent localization of camouflaged objects. B–D. Improvement in behavioral accuracy when objects move compared to when they remain static, plotted as a function of static-image error for horizontal position (B), vertical position (C), and object size (D). Motion produces a clear improvement for object position estimates, particularly for stimuli that are difficult under static viewing conditions. The effect is weaker for size estimates, consistent with the lower reliability of human size judgments. Errorbars denote standard deviation across sampling over repetitions to perform the analysis in a cross-validated way.

Together, these results demonstrate that motion provides a robust cue for estimating object attributes in naturalistic scenes. Importantly, the benefit of motion was strongest for stimuli that were difficult to interpret from static images, suggesting that temporal information helps disambiguate objects that are otherwise visually indistinguishable from their background.

### 4.2 Comparison of image and video ANNs on decoding motion-based object attributes under camouflage

Before testing whether models benefit from motion, it is first necessary to establish that their internal representations contain information about the object attributes measured in our behavioral task. If model features encode object position, size, or motion information, these attributes should be recoverable using simple linear decoding. In all following analyses, accuracy was quantified as the Spearman correlation between the decoded object attribute and the corresponding ground-truth annotation across stimuli. This correlation-based metric measures how well variations in the predicted attribute track variations in the true object attribute across images. Higher correlations, therefore indicate more accurate representations of the object property.

We observe that all the considered models are able to decode object attributes from the moving object video stimuli. Image-based models processed individual frames independently, while video-based models integrated information across time using spatiotemporal filters learned during their respective optimizations. We observed that both image-based and video-based ANNs achieve above-chance decoding accuracy for object properties such as horizontal and vertical position and size (Fig. 6 A–C). However, a clear gap remains between ANNs and human performance (also estimated here as a correlation with ground truth), with humans consistently achieving the highest accuracy across object position attributes. Estimating object size results in a more challenging task for humans, which is consistent with an overall reduced human reliability (Fig. 2 E). Surprisingly, however, in this task, humans are surpassed by several image-based and video-based models. Interestingly, despite the architectural differences between model classes, image-based and video-based neural networks showed similar decoding accuracy for object position and size. Because these attributes can be inferred from spatial structure within individual frames, temporal integration does not provide a substantial advantage for these tasks (from an accuracy point of view).

**Fig. 6.**
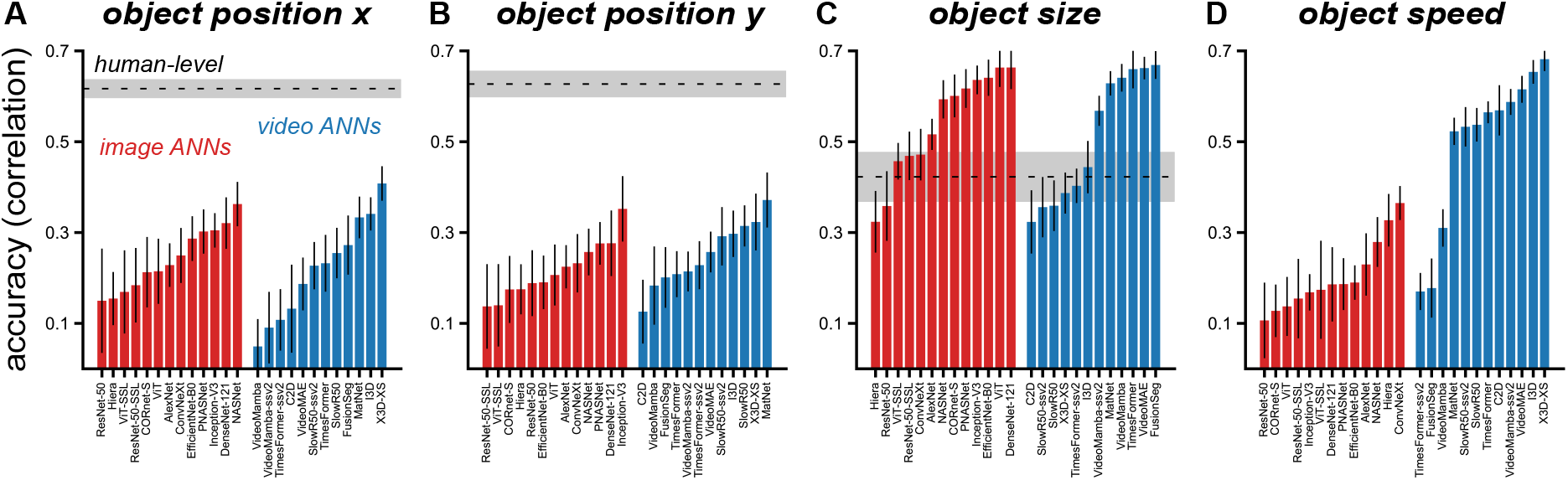
Humans and artificial neural networks differ in estimating object attributes from camouflaged scenes. Decoding accuracy (Spearman correlation with ground truth) for object size (A), horizontal position (B), vertical position (C), and object speed (D). Both image-based and video-based models allow reliable decoding of spatial object attributes such as position and size, indicating that these form attributes are represented in model features. However, humans outperform artificial models for estimating object position. In contrast, models outperform humans for object size estimation, consistent with the lower reliability of human size judgments. For the dynamic attribute (object speed), video-based architectures substantially outperform image-based models, reflecting their ability to integrate temporal information across frames.

A different pattern emerged for object speed (Fig. 6 D), which represents a dynamically defined attribute that requires integrating information across time. On this task, video-based neural networks substantially outperformed image-based models (Wilcoxon rank-sum *p <* 0.001). This advantage is expected because video architectures explicitly integrate information across frames using spatiotemporal operations, whereas image-based models process each frame independently.

These findings establish that object form attributes are represented in artificial neural networks and therefore provide a foundation for testing whether models exhibit the same motion-dependent improvements observed in human perception.

### 4.3 Video-based models outperform image-based models in motion exploitation

As demonstrated earlier, human observers show improved performance when objects move. If ANNs capture a similar computation, models that integrate temporal information should benefit from motion in ways that frame-based models cannot. We therefore next evaluated whether ANNs capture the same benefits of motion observed in human behavior.

Decoding analyses revealed a clear performance difference between image-based and video-based models (Fig. 7 D-F). Image based ANNs exhibited no significant improvement when predicting object form attributes from dynamic stimuli relative to static frames (*x*-position, Fig. 7 A, D: mean Δ accuracy = 0.0 *±*0.011, *p* = 0.96, *t*(12) = *−*0.07; *y*-position, Fig. 7 B, E: mean Δ accuracy = *−*0.02*±* 0.010, *p* = 0.090, *t*(12) =*−* 1.85; size, Fig. 7 C, F: mean Δ accuracy = 0.01*±* 0.0087, *p* = 0.05, *t*(12) = 2.14). In contrast, video-based architectures showed significantly improved accuracies when motion cues were present (*x*-position: mean Δ accuracy = 0.06*±* 0.021, *p* = 0.016, *t*(11) = 2.83; *y*-position: mean Δ accuracy = 0.07*±* 0.017, *p* = 0.004, *t*(11) = 3.65; size: mean Δ accuracy = 0.09 *±*0.017, *p <* 0.001, *t*(11) = 5.07).

**Fig. 7.**
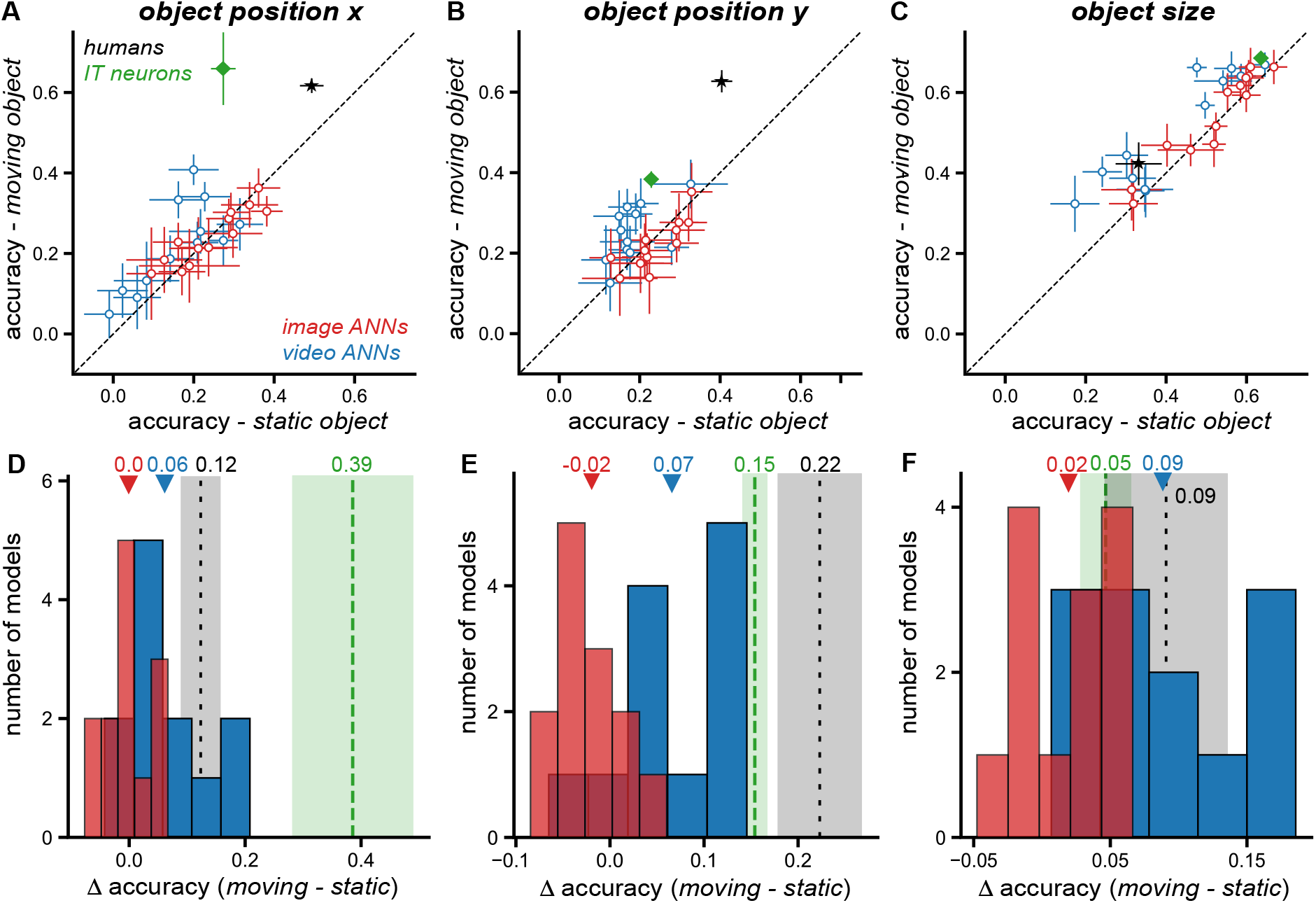
Video-based models exploit motion cues more effectively than image-based models. (A–C) Decoding accuracy (Spearman correlation with ground truth) for object horizontal position (A), vertical position (B), and object size (C) when models are evaluated on static-object frames versus moving-object videos. Each point represents a model, grouped by architecture class (image-based in red, video-based in blue). Neural decoding results from macaque IT cortex (green diamond) and human behavioral performance (black star) are shown for reference.(D–F) Motion-dependent improvement in decoding accuracy, defined as the difference between dynamic and static conditions (moving*−* static), for horizontal position (D), vertical position (E), and object size (F). Histograms show the distribution of improvements across model families. Video-based architectures exhibit significantly larger motion-dependent improvements than image-based models for horizontal position (video ANN mean: 0.06*±* 0.021 SE; image ANN mean: 0.0*±* 0.011 SE; Wilcoxon rank-sum *p* = 0.03), vertical position (video ANN mean: 0.07 *±* 0.017 SE; image ANN mean: *−* 0.02*±* 0.010 SE; *p* = 0.001), and object size (video ANN mean: 0.09*±* 0.017 SE; image ANN mean: 0.02*±* 0.0087 SE; *p* = 0.002). Together, these results demonstrate that models incorporating temporal processing capture the motion-dependent advantages observed in human vision (and visible in non human primate brain mechanisms) more effectively than frame-based architectures.

These findings demonstrate that temporal integration improves model performance on tasks involving camouflaged objects, supporting the idea that motion provides a useful signal for stabilizing object representations. Importantly, although this improvement does not fully match the magnitude of the motion advantage observed in humans, video-based ANNs show improvement closer in magnitude to those observed in humans, compared to image ANNs. These results suggest that video-based models provide a more biologically plausible account of human behavior in dynamic visual environments than image-based models.

### 4.4 Video models exhibit a gap with human behavioral patterns

To evaluate whether artificial neural networks capture the perceptual strategies used by humans, we compared model predictions directly with human behavioral responses across stimuli. Importantly, a model may achieve high decoding accuracy while still relying on visual cues that differ from those used by human observers. To assess this possibility, we quantified the consistency between model-derived predictions and human behavioral estimates (Fig. 8 A–C) across stimuli (at a video-by-video grain). This analysis evaluates whether models reproduce the stimulus-by-stimulus pattern of human perceptual judgments rather than simply achieving high overall accuracy.

**Fig. 8.**
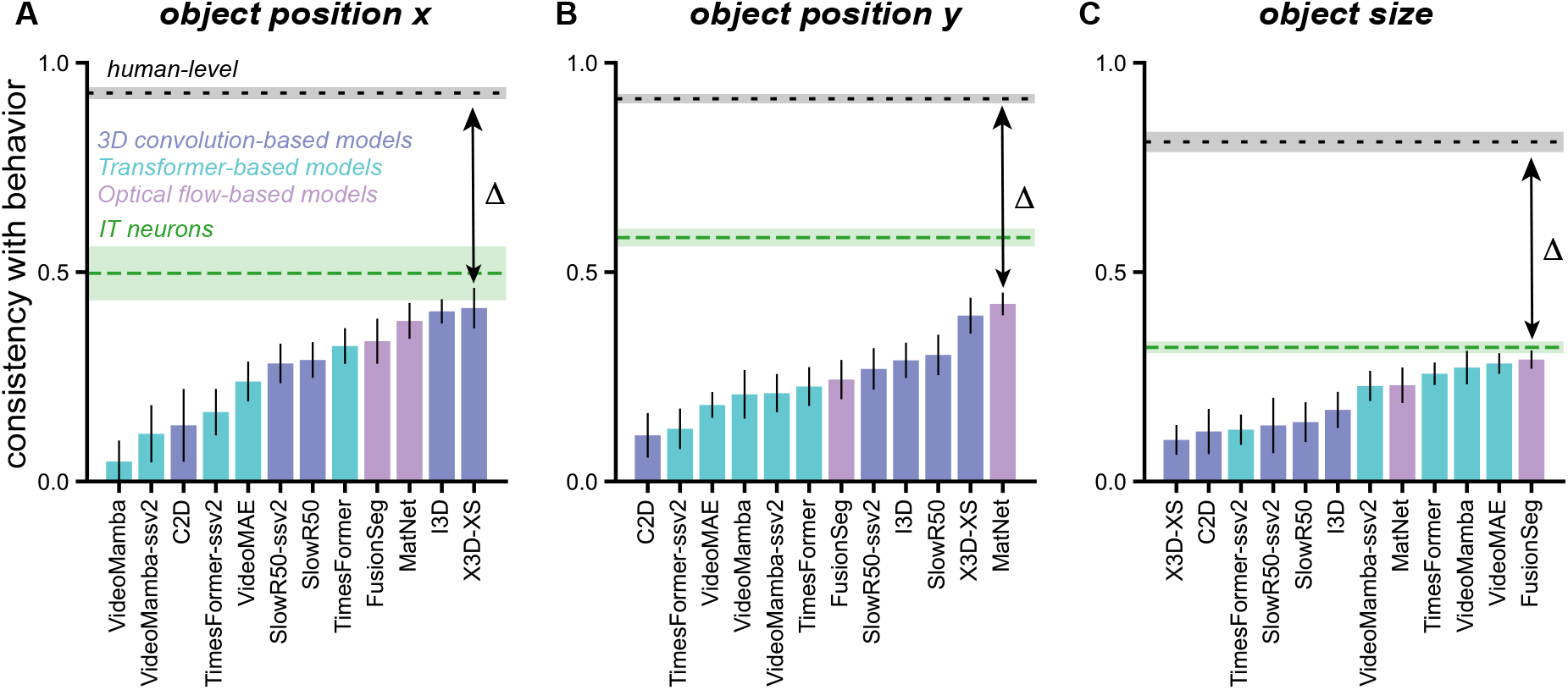
Comparison of human behavioral patterns, IT decoding predictions, and video-based models. A–C. Consistency between model predictions and human behavioral patterns for estimating horizontal object position (A), vertical object position (B), and object size (C). Consistency is measured as the correlation between model-derived decoding patterns and human behavioral responses across stimuli. Results are shown for different families of video-based architectures, including 3D convolution-based, transformer-based, and optical flow–based models, and are compared with predictions obtained from IT neural responses. The black dashed line indicates the human-level ceiling, estimated from trial split half consistency. Overall, IT neural activity shows the highest alignment with human behavior, while among artificial models, 3D convolution-based architectures and optical flow–based methods achieve higher consistency than transformer models, although all models remain below the human benchmark (Δ).

Overall, video ANNs showed significant (spearman correlations *>* 0, *p <* 0.05) alignment with human behavioral patterns, but their consistency remained significantly below the human-level ceiling estimated from repetition split-half reliability. We found two distinct patterns in these comparisons. First, the strongest and most interpretable model–human alignment emerged for object position (both horizontal and vertical). Several video models showed moderate consistency with human responses, indicating that their predictions captured aspects of the stimulus-by-stimulus variability in human localization judgments. In contrast, object size estimates showed a significantly lower model–human correspondence across architectures. This likely reflects the lower reliability of human size judgments in this task, which reduces the measurable ceiling for model–human alignment and makes it harder to distinguish differences between models.

Second, within the object position results, we observed systematic differences across model architectures. Convolution-based video models and architectures incorporating explicit motion signals (e.g., optical-flow-based methods) generally showed higher behavioral consistency than transformer-based video models (*x*-position, *y*-position, Wilcoxon ranksum *p* = 0.04). One possible interpretation is that position estimation in camouflaged scenes depends strongly on preserving local motion-defined spatial structure, such as motion boundaries and the spatial distribution of foreground signals. Architectures based on convolutional spatiotemporal filtering may therefore better capture the cues used by human observers to localize objects in motion-defined scenes. In contrast, the pattern for object size differed across architectures: transformer-based video models showed higher behavioral consistency with human size estimates than convolution-based models. This suggests that estimating object extent may rely more heavily on integrating information across larger spatial regions of the scene, a computation that may benefit from the global aggregation mechanisms characteristic of transformer architectures.

Overall, our evidence suggests that while modern video models capture aspects of the computations underlying human perception of object form under camouflage, important representational differences might remain between artificial and biological visual systems.

### 4.5 Macaque IT cortex encodes object attributes more accurately during motion

Previous work has shown that humans and macaques exhibit strong behavioral alignment on object recognition tasks under both static (Rajalingham et al. 2018) and dynamic (Ramezanpour et al. 2024) viewing conditions. In addition, neural population activity in macaque inferior temporal (IT) cortex has been shown to contain representations that support and explain these behavioral judgments (Kar and DiCarlo 2024; Hong et al. 2016; Hung et al. 2005; Majaj et al. 2015). Because IT lies at the apex of the ventral visual stream and plays a key role in object perception, it provides a useful biological reference for interpreting the relationship between human behavior and artificial neural network models.

We therefore asked whether the behavioral benefits of motion observed in humans are also reflected in neural representations within IT cortex. If motion improves object perception by stabilizing object representations, then neural population activity in IT should contain more reliable information about object attributes when objects move compared to when they remain static.

We recorded neural activity from the IT cortex of two monkeys while macaque monkeys passively viewed the same set of video stimuli shown to the human participants. We then trained linear decoders to estimate object horizontal position, vertical position, and object size from IT population activity.

Population decoding analyses revealed that object form attributes could be reliably predicted from IT neural activity. Linear decoders trained on neural population responses successfully estimated object horizontal position, vertical position, and object size (Fig. 7 A-C green dots). Crucially, decoding accuracy was higher for moving object stimuli than for static object frames. For example, decoding object size from neural responses yielded stronger correlations with ground-truth values when animals viewed dynamic videos compared to static images (position *x* Δ = 0.05, *p <* 0.001, permutation test). Similar improvements were observed for both horizontal position (Δ*error* = 0.39, *p <* 0.001, permutation test) and vertical position estimates (size Δ = 0.15, *p <* 0.001, permutation test). Consistent with video ANNs, and previous results (Ramezanpour et al. 2024), we also observed that IT responses significantly predicted object speed with an accuracy (correlation with ground truth) of 0.5.

These results indicate that motion enhances the fidelity of object form attribute representations in IT cortex. Rather than representing objects purely based on static appearances, ventral visual populations appear to incorporate temporal information that improves the stability of object representations under challenging visual conditions.

### 4.6 Video models partially align with IT decoding yet fall short of human behavioral alignment

We next asked how closely video ANNs reproduce the neural decoding patterns observed in macaque IT cortex and whether such alignment explains their ability to predict human behavioral responses. If IT representations provide the neural substrate underlying perceptual judgments of object form attributes, then models whose internal representations more closely resemble IT activity should better reproduce human behavioral patterns.

To test this prediction, we first compared model decoding performance directly with neural decoding results obtained from IT cortex. For each model, we computed correlations between predicted and ground-truth object attributes for the dynamic stimulus conditions. When neural decoding performance was plotted alongside model results (Fig. 8), IT population responses aligned most closely with the best-performing video models. In particular, several video architectures achieved decoding correlations comparable to neural decoding performance for object position and size estimation. This suggests that temporal integration in some video-based models enables them to approximate aspects of the motion-dependent computations observed in ventral visual cortex (as previously shown in Dunnhofer et al. (2026a)). Importantly, this comparison provides a conservative benchmark for model performance. Neural decoding results are based on finite samples of recorded neurons, whereas artificial models operate with much larger representational populations. We observed that neural–behavior alignment improves as the number of sampled neurons increases, suggesting that larger neural populations would likely provide even stronger behavioral predictions. Therefore, important differences remained. While specific video models captured some of the improvements associated with motion, they did not fully reproduce the magnitude or pattern of behavioral and neural advantages observed in biological vision.

To further investigate why some video ANNs align better with humans, we examined whether the similarity between ANN representations and neural responses predicted the degree to which models captured human behavioral patterns. We quantified the representational alignment between each model and IT using Centered Kernel Alignment (CKA) and compared these scores with each model’s consistency with human behavior (Fig. 9). Across models, higher CKA similarity to IT was associated with greater consistency with human behavioral responses for object position decoding (*x*: *r* = 0.75, *p* = 0.005; *y*: *r* = 0.70, *p* = 0.011), indicating that models whose internal representations more closely resemble IT activity tend to better predict human perceptual judgments. Interestingly, this relationship was specific to object position. For object size, neural alignment did not predict behavioral consistency (Fig. 9 C; *r* =*−*0.05, *p* = 0.88).

**Fig. 9.**
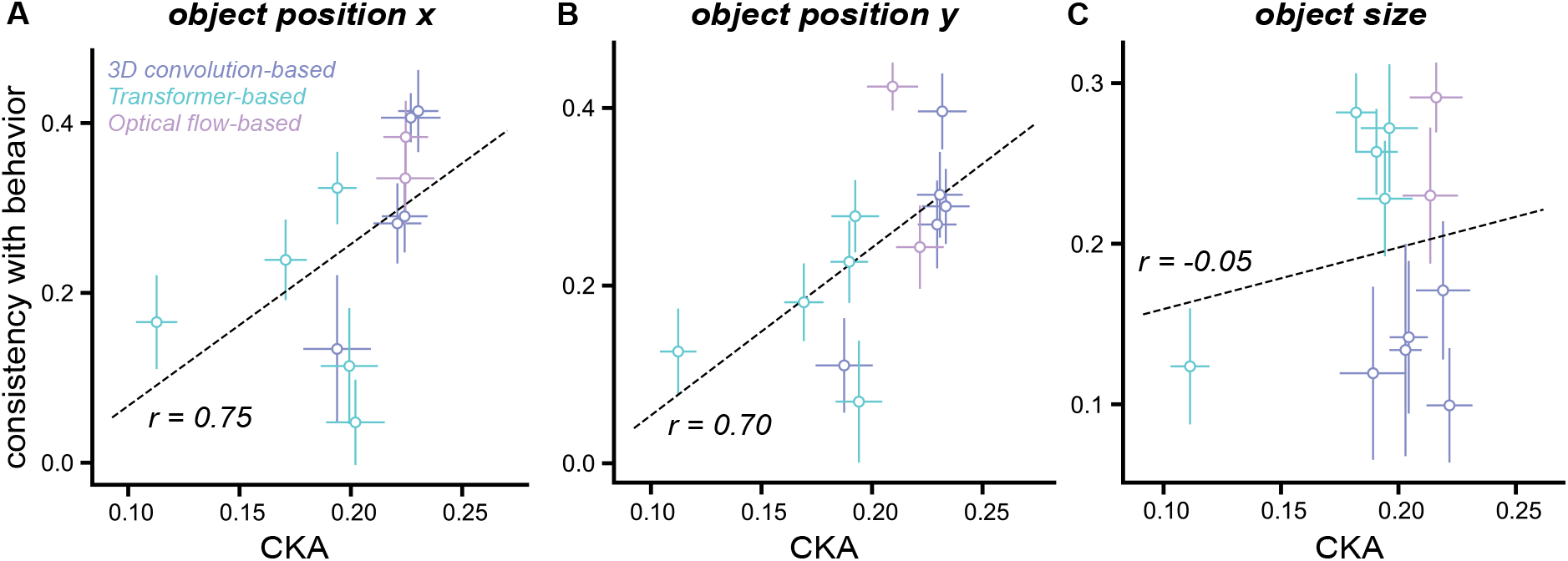
Neural alignment predicts model consistency with human behavior. A–C. Relationship between ANN–IT representational similarity, measured using CKA, and consistency of the ANN with human behavior for decoding horizontal object position (A), vertical object position (B), and object size (C). Each point represents a model, grouped by architecture family (3D convolution-based, transformer-based, and optical flow–based models). For position decoding, higher CKA scores are associated with greater consistency with human behavior, indicating that models whose representations more closely match IT activity better capture human perceptual patterns. 3D convolution-based and optical flow–based models show the highest levels of neural and behavioral alignment, while transformer-based models exhibit lower correspondence. Despite this relationship, overall CKA values remain relatively low, highlighting the gap between current model representations and primate IT cortex.

Comparing model families revealed systematic differences. 3D convolution-based models and optical-flow-based architectures achieved the highest levels of both neural alignment and behavioral consistency, whereas transformer-based models generally showed lower correspondence with IT responses and weaker agreement with human behavior. Despite this positive relationship between neural similarity and behavioral predictivity, the overall magnitude of the CKA scores remained relatively low across models (Fig. 9 A–C), suggesting that current architectures still diverge substantially from the representational geometry observed in primate IT cortex.

Taken together, these results indicate that although modern video-based ANNs capture aspects of the neural computations underlying motion-based object perception, their representations remain insufficient to fully explain human perceptual behavior. Models that more closely match IT representations better predict human behavior, but important representational differences remain between artificial and biological visual systems.

## 5 Discussion

Our results provide converging evidence that motion plays a critical role in stabilizing object perception in challenging visual environments. Across human behavior, neural population responses in macaque inferior temporal (IT) cortex, and video-based artificial neural networks, dynamic stimuli containing moving objects improved the estimation of object attributes such as position and size compared with static object frames. These findings suggest that motion is not merely an auxiliary signal for action recognition or object tracking, but can directly support the computation of object form when appearance cues are unreliable.

From a computer vision perspective, our results highlight an important limitation of current image-based recognition systems (Krizhevsky et al. 2012; He et al. 2016; Dosovitskiy et al. 2020). Models that process frames independently showed little benefit from motion information even when dynamic cues were highly informative. In contrast, video-based neural networks that integrate temporal information achieved significantly better performance on these tasks. This result supports the growing view that temporal integration is an important ingredient for robust visual recognition in dynamic environments (Ramezanpour et al. 2024; Dunnhofer et al. 2026a).

However, even the best-performing video models (Feichtenhofer 2020; Bertasius et al. 2021; Tong et al. 2022; Li et al. 2024) did not fully reproduce the behavioral and neural advantages observed in biological systems. Although several architectures showed improved performance on dynamic stimuli, their improvements were typically smaller or less consistent than those observed in humans and primate neural populations. This gap suggests that current video architectures may not yet capture the computational mechanisms used by biological vision to exploit motion cues for object perception.

One possible explanation is that biological systems integrate motion and form information more tightly than current artificial models. In many video architectures (Feichtenhofer 2020; Bertasius et al. 2021; Tong et al. 2022; Li et al. 2024; Zhou et al. 2020), temporal processing is primarily optimized for recognizing actions or temporal events rather than stabilizing spatial representations of objects. By contrast, the primate ventral visual stream may combine temporal and spatial signals in a way that directly enhances the reliability of object representations. In such a framework, motion signals may help disambiguate object boundaries, stabilize object localization, and reinforce form representations when static appearance cues are degraded.

More broadly, our findings contribute to a growing body of work suggesting that dynamic visual processing plays an important role in ventral stream computations (Bigelow et al. 2023; Ramezanpour et al. 2024; Dunnhofer et al. 2026b). While classical models of the ventral visual stream emphasize static form processing, our results indicate that temporal information can significantly enhance the fidelity of ventral representations when appearance cues are unreliable. Motion-based form computation may therefore represent an important principle shared by both biological and artificial vision systems.

### 5.1 Limitations

Several limitations of the present study should be considered. First, our experiments focused on a specific class of naturalistic stimuli involving camouflaged animals. While these scenes provide a useful testbed for studying motion-based perception, additional work will be needed to determine whether similar motion advantages generalize to other challenging visual conditions, such as objects embedded in cluttered scenes, partially occluded objects, or objects undergoing viewpoint and illumination changes. Second, although we evaluated a broad range of artificial neural network architectures, the space of possible video models is evolving rapidly. Future work may identify architectures that more effectively capture the temporal computations observed in biological vision. In particular, models that more tightly integrate spatial and temporal representations may provide improved alignment with neural and behavioral data. Third, our neural analyses focused on decoding object attributes from population activity in IT cortex. While decoding analyses reveal the information available in neural representations, they do not directly identify the circuit mechanisms that generate those representations. Understanding how motion information is integrated across visual areas, including interactions between dorsal and ventral streams, remains an important direction for future research. Finally, our comparison between biological and artificial systems relied primarily on linear decoding analyses and representational similarity measures such as CKA. These approaches provide standardized and interpretable benchmarks for model–brain comparison while minimizing assumptions about the form of the mapping between systems. However, richer analyses of representational structure—such as examining temporal population trajectories, motion-specific subspaces, or the geometry of neural manifolds—may provide deeper insight into the computations underlying motion-based object perception.

## 6 Conclusion

We introduced a set of benchmarks to investigate how motion contributes to object perception in challenging visual environments. Using videos of camouflaged animals, we compared human behavior, neural population responses in macaque IT cortex, and artificial neural networks on the task of estimating object position and size. Our results show that humans benefit systematically from motion when estimating object attributes. Imagebased neural networks achieve strong static accuracy but do not exhibit this motion-dependent improvement. Several video-based architectures reproduce this behavioral pattern, and the degree to which they do so is predicted by their representational similarity to IT population responses.

These findings suggest that static object accuracy alone is insufficient to evaluate models of visual perception. Instead, models should also capture the motion-dependent computations that support robust perception in biological vision. More broadly, our results highlight the importance of using biological representations as reference systems for guiding the development of artificial models that better capture the dynamic computations underlying natural vision.

